# Early evolution of beetles regulated by the end-Permian deforestation

**DOI:** 10.1101/2021.10.12.464043

**Authors:** Xianye Zhao, Yilun Yu, Matthew E. Clapham, Evgeny Yan, Jun Chen, Edmund A. Jarzembowski, Xiangdong Zhao, Bo Wang

**Affiliations:** State Key Laboratory of Palaeobiology and Stratigraphy, Nanjing Institute of Geology and Palaeontology and Center for Excellence in Life and Paleoenvironment, Chinese Academy of Sciences, Nanjing 210008, China; University of Chinese Academy of Sciences, Beijing 100049, China; Institute of Vertebrate Paleontology and Paleoanthropology, Chinese Academy of Sciences, Beijing 100044, China; Department of Earth and Planetary Sciences, University of California, Santa Cruz, CA 95064, USA; Palaeontological Institute, Russian Academy of Sciences, ul. Profsoyuznaya 123, Moscow, 117647 Russia.; Institute of Geology and Paleontology, Linyi University, Linyi 276000, China; Department of Earth Sciences, Natural History Museum, London, SW7 5BD, UK

## Abstract

The end-Permian mass extinction (EPME) led to a severe terrestrial ecosystem collapse. However, the ecological response of insects—the most diverse group of organisms on Earth—to the EPME remains poorly understood. Here, we analyse beetle evolutionary history based on taxonomic diversity, morphological disparity, phylogeny, and ecological shifts from the Early Permian to Middle Triassic, using a comprehensive new data set. Permian beetles were dominated by xylophagous stem groups with a high diversity and disparity, which probably played an underappreciated role in the Permian carbon cycle. Our suite of analyses shows that Permian xylophagous beetles suffered a severe extinction during the EPME largely due to the collapse of forest ecosystems, resulting in an Early Triassic gap of xylophagous beetles. New xylophagous beetles appeared widely in the early Middle Triassic, which is consistent with the restoration of forest ecosystems. Our results highlight the ecological significance of insects in deep-time terrestrial ecosystems.

## INTRODUCTION

The end-Permian mass extinction (EPME; approximately 252 million years ago) was the most severe extinction event in the Phanerozoic (*Benton and Newell, 2014*). The EPME was primarily caused by the eruption of the Siberian flood basalts (*Burges and Bowring, 2015*; Fielding et al., 2019), which generated excessive emissions of thermogenic methane, CO_2_ and SO_2_ that cascaded rapid global warming (Wu et al., 2021; Black et al., 2020), oceanic acidification and anoxia/euxinia (Schobben et al., 2020), aridification and other shifts in the hydrological cycle (Sun et al., 2012), acid rain (Black et al., 2014), wildfires (Shen et al., 2011), and ozone destruction (Benca et al., 2018). The response of terrestrial ecosystems to the EPME is quite heterogeneous probably due to biotic, topographic and latitudinal differences (Fielding et al., 2019; Zhao et al., 2020; *Dal Corso et al., 2020*). Moreover, how terrestrial ecosystems were affected during the EPME is still highly controversial (*Benton and Newell, 2014*; Gastaldo, 2019). Terrestrial tetrapods and plants are considered to have been severely affected by the EPME mostly based on diversity and taxonomic composition (*Benton and Newell, 2014*; Viglietti et al., 2021); however, such mass extinction was questioned by a more comprehensive dataset of plant macro- and microfossils (Gastaldo, 2019; Nowak et al., 2019). Similarly, Permian insects are thought to have suffered a significant extinction (*Labandeira and Sepkoski, 1993*; *Béthoux et al., 2005; Labandeira, 2005*; Condamine et al., 2020; Condamine et al., 2016), but this was not supported by other molecular phylogenetic and fossil record analyses (Ponomarenko, 2016; Dmitriev et al., 2018; Montagna et al., 2019; Schachat et al., 2019). In addition, the ecological response of insects to the EPME remains poorly understood (*Benton and Newell, 2014*; *Schachat and Labandeira, 2020*).

Beetles (Coleoptera) are the most speciose group of extant insects (Stork, 2018), with a stratigraphic range dating back to at least the lowest Permian (Ponomarenko, 2016; Kirejtshuk et al., 2014). They have a rich fossil record since the Permian and display a wide array of lifestyles (*Figure 1*) (*Ponomarenko, 1969, 2003*). Their fossil record thus offers a unique and complementary perspective for studying the ecological response of insects to the EPME. The evolutionary history of Coleoptera has been widely investigated through molecular phylogenetic analyses (Condamine et al., 2016; *McKenna et al.*, 2019; Zhang et al., 2018), morphological phylogenetic analyses (Beutel et al., 2008; Beutel et al., 2019), and fossil record analyses (*Ponomarenko, 2003, 2016*; *Smith and Marcot, 2015*). Although a long-term Palaeozoic-Mesozoic turnover of beetle assemblages is supported by almost all analyses, the detailed ecological response to the EPME and its explanatory mechanisms remain unclear. Most of the Permian and Triassic beetles belong to stem groups (extinct suborders or families; *Figure 1*), and thus they show character combinations and evolutionary history that cannot be inferred or predicted from phylogenetic analysis of modern beetles. In particular, two problems were ignored by previous analyses. First, phylogenetic relationships of some key fossils remain poorly resolved, particularly in their evolutionary relationships to modern taxa. Second, there are two complementary taxonomic systems for Permian and Triassic beetles: one is artificial formal taxa (based on isolated elytra that cannot be definitely classified into any natural group), and the other is the natural taxonomy (commonly based on more complete fossils including bodies and elytra). The formal taxa, like trace fossil taxa, lack comprehensive phylogenetic data (Ponomarenko, 2004), and thus they cannot be used unreservedly for biodiversity and phylogenetic analyses, but can be helpful in the morphospace analysis. These issues cloud the temporal resolution of coleopteran biodiversity in deep time and complicate the evolutionary trajectory of beetles but can be overcome through a combination of multiple analytical methods. Therefore, taxonomic diversity, morphological disparity, and ecological shifts are best evaluated jointly to better understand how the EPME has shaped the evolutionary history of beetles.

**Fig. 1.**
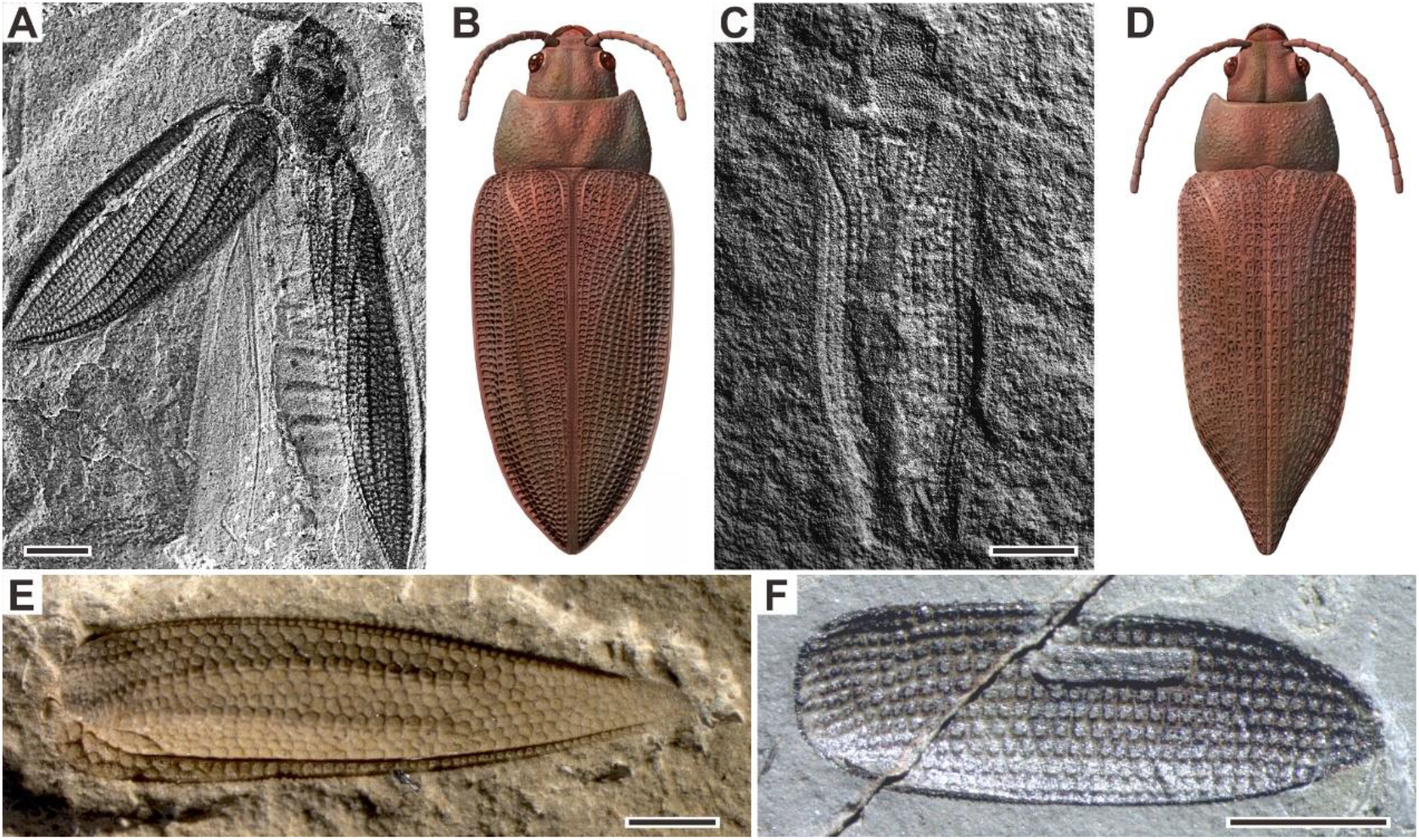
Examples of Permian beetles. (**A** and **B**) Tshekardocoleidae, *Moravocoleus permianus* Kukalova, 1969, photograph and reconstruction. (**C** and **D**) Permocupedinae, *Permocupes sojanensis* Ponomarenko, 1963, photograph and reconstruction. (**E**) Tshekardocoleidae, *Sylvacoleus richteri* Ponomarenko, 1963, elytra photograph. (**F**) Taldycupedinae, *Taldycupes reticulatus* Ponomarenko, 1969, elytra photograph. Scale bars represent 1 mm.

Here, we compile an updated database of beetles from the Early Permian to Middle Triassic based on the taxonomic revision of fossils (including formal taxa). We analyse the evolution of taxonomic diversity, morphological disparity and palaeoecological shifts of beetles from the Early Permian to Middle Triassic through phylogenetic and palaeoecological reconstructions and morphospace analyses of fossil material. Our results suggest that xylophagous (wood feeding) beetles probably played a key and underappreciated role in the Permian carbon cycle and that the EPME had a profound ecological influence on beetle evolution. These results provide new insights into the ecological role of insects in deep-time terrestrial ecosystems and the ecological response of insects to deforestation and global warming.

## RESULTS

### Taxonomic diversity

We compiled an updated database of beetles (22 families, 125 genera, and 299 species) from the Early Permian to Middle Triassic based on the taxonomic revision of natural and formal taxa (*Supplementary Dataset S1*)). Our database contains 19 families, 109 genera and 220 species of natural taxa. There is a steady increase of families from the Early Permian to Middle Triassic, which is consistent with the result of Smith and Marcot (*Smith and Marcot, 2015*) whose analyses were only conducted at the family level. The diversity of natural taxa displays almost the same trajectory at both species and genus levels (*Figure 2C, E*). The diversity is roughly stable in the Early Permian (Cisuralian), mainly represented by Tshekardocoleidae (*Figure 1*), increases rapidly in the Middle Permian (Guadalupian) and Late Permian (Lopingian), with the rise of the major clades Permocupedidae (Permocupedinae and Taldycupedinae) and Rhombocoleidae. Subsequently, it plunges in the Early Triassic and recovered gradually from the Anisian (early Middle Triassic). In the Ladinian (late Middle Triassic) the diversity clearly exceeds that of the Late Permian (*Figure 2C, E*). From the Middle Triassic, the Permian coleopteran assemblage characterised by Tshekardocoleidae, Permocupedidae and Rhombocoleidae is completely replaced by a Triassic assemblage dominated by Cupedidae, Phoroschizidae and Triaplidae.

**Fig. 2.**
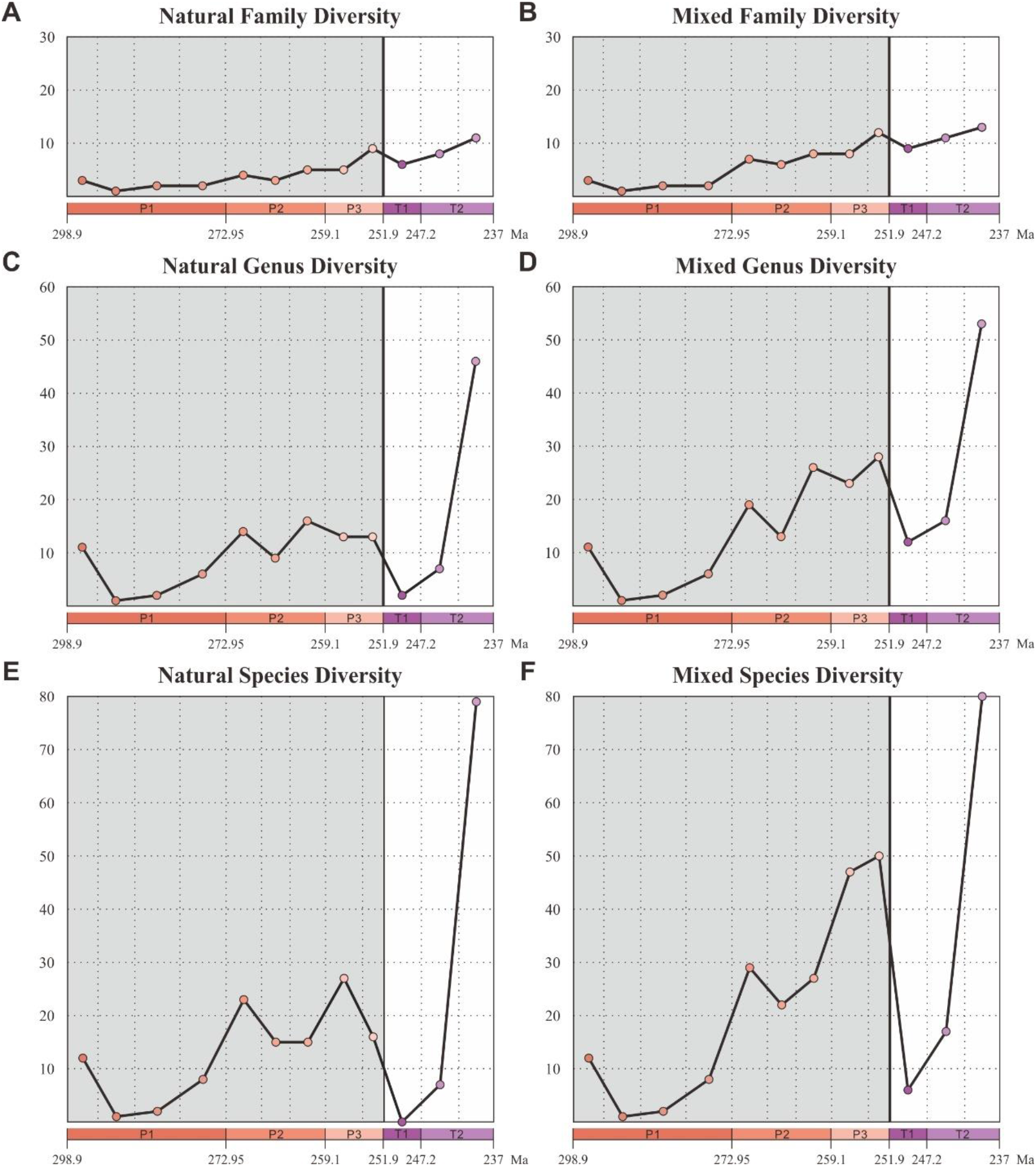
Diversity of Coleoptera from the Early Permian to Middle Triassic. Natural taxa and mixed taxa (natural taxa and formal taxa) are counted at family, genus, and species levels separately. (**A**) family-level diversity of natural taxa. (**B**) family-level diversity of mixed taxa. (**C**) genus-level diversity of natural taxa. (**D**) genus-level diversity of mixed taxa. (**E**) species-level diversity of natural taxa. (**F**) species-level diversity of mixed taxa. P1, Early Permian; P2, Middle Permian; P3, Late Permian; T1, Early Triassic; T2, Middle Triassic.

Our database also contains three families, 17 genera and 79 species of formal taxa. A considerable proportion of Permian beetles belong to such taxa (Permosynidae, Schizocoleidae, and Asiocoleidae). These formal taxa mostly belong to stem groups, but some should probably be attributed to the two extant suborders Adephaga and Polyphaga. Both species and genus-level diversities of formal taxa show a gradual increase from the Middle to Late Permian, but decrease distinctly from the Triassic (*Appendix 1—figure 1*). The mixed taxa diversity (combining natural and formal taxa) displays the same trajectory to that of natural taxa at both species and genus levels (*Figure 2D, F*).

### Phylogeny

We carried out a phylogenetic analysis based on 93 adult and larval characters across 15 natural taxa representing all natural families of Coleoptera from the Early Permian to Middle Triassic (*Supplementary Dataset S2*). Our parsimony analysis result is consistent with a previous analysis (Beutel et al., 2008), and confirms that Tshekardocoleidae, Permocupedidae (Permocupedinae and Taldycupedinae), Rhombocolediae, and Triadocupedinae are the stem group of Coleoptera (*Figure 3A* and *Appendix 1—figure 2*).

**Fig. 3.**
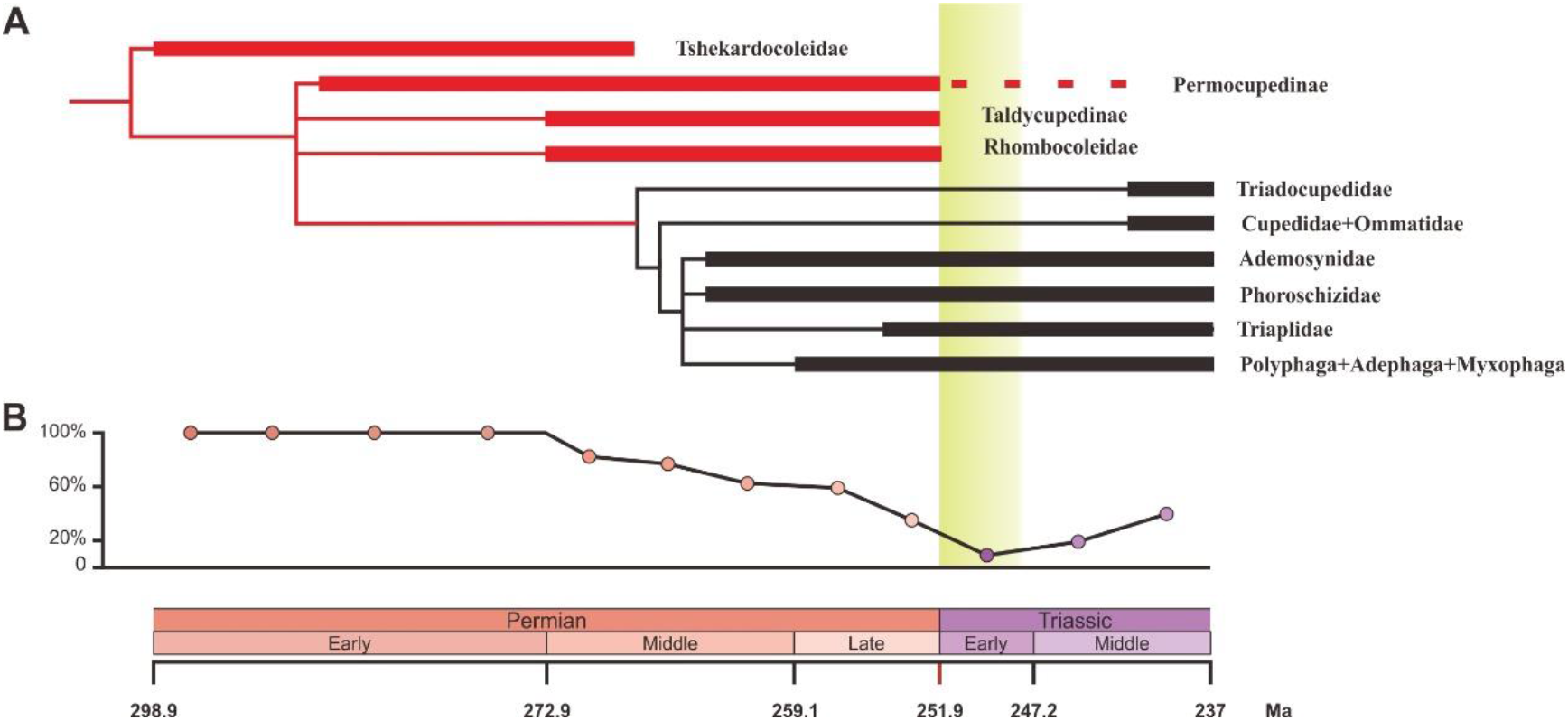
Ecological shifts of Coleoptera from the Early Permian to Middle Triassic. (**A**) Simplified phylogeny of Coleoptera from the Early Permian to Middle Triassic. Thick lines indicate the known extent of the fossil record. The branches representing stem groups are shown in red. The “dead clade walking” pattern is symbolized by the dashed line. Details of the phylogenetic analysis see *Appendix 1—figure 2* and *Appendix, Supplemental Text A, B and D*. (**B**) Genus percentage of xylophagous groups from the Early Permian to Middle Triassic. Yellow graded band represents the ‘coal gap’.

### Morphological disparity

We chose the beetle elytra—hardened forewings primarily serving as protective covers for the hindwings and body underneath—to perform the morphological disparity analysis for three reasons: 1) elytra are the most commonly preserved fossils of Palaeozoic and Mesozoic beetles, and they are easily accessible in the literature and in online databases; 2) Permian and Triassic elytra display complex morphological structure (Ponomarenko, 1969); 3) elytra morphology has long been studied in relation to taxonomic diversity of living and extinct beetles (Ponomarenko, 2004; Tong et al., 2021).

We assembled two discrete character matrices (at species and genus levels) based on 35 characters of 197 genera and 346 species (including undetermined species and unnamed specimens) for morphological disparity analyses *(Supplementary Dataset S3*) (Lloyd, 2016). The taxa were ordinated into a multivariate morphospace using both principal coordinates analysis (PcoA) and non-metric multidimensional scaling (NMDS) with two distance metrics, including the Generalized Euclidean Distance (GED) and maximum observable rescale distance (MORD). We chose both sum of variance (sov) and product of variance (pov) as the proxy for morphological disparity due to their robustness in sample size (*Simões et al., 2020*). The use of discrete characters produces results that have nonmetric properties, but this approach can be used to elucidate broad patterns of similarities and clustering within multidimensional space (Deline et al., 2018).

The patterns of morphospace occupation of beetles in different time-bins are shown in three-dimensional plots delimited by combinations of the first three axes of the PcoA and NMDS results based on MORD metrics (*Figure 4A* and *Appendix—figure 3A–C*). The morphological disparity results of two ordination methods within the MORD and GED matrix shows the same trajectory at both genus and species levels. The evolutionary pattern of morphological disparity is robust in different disparity metrics. The disparity is low in the Early Permian, with a significant increase during the Middle Permian. It is roughly stable in the Middle and Late Permian, subsequently showing a distinct plunge in the Early Triassic, slightly recovering in the Middle Triassic but is still significantly lower than in the Middle and Late Permian (*Figure 4* and *Appendix—figures 3–10*).

**Fig. 4.**
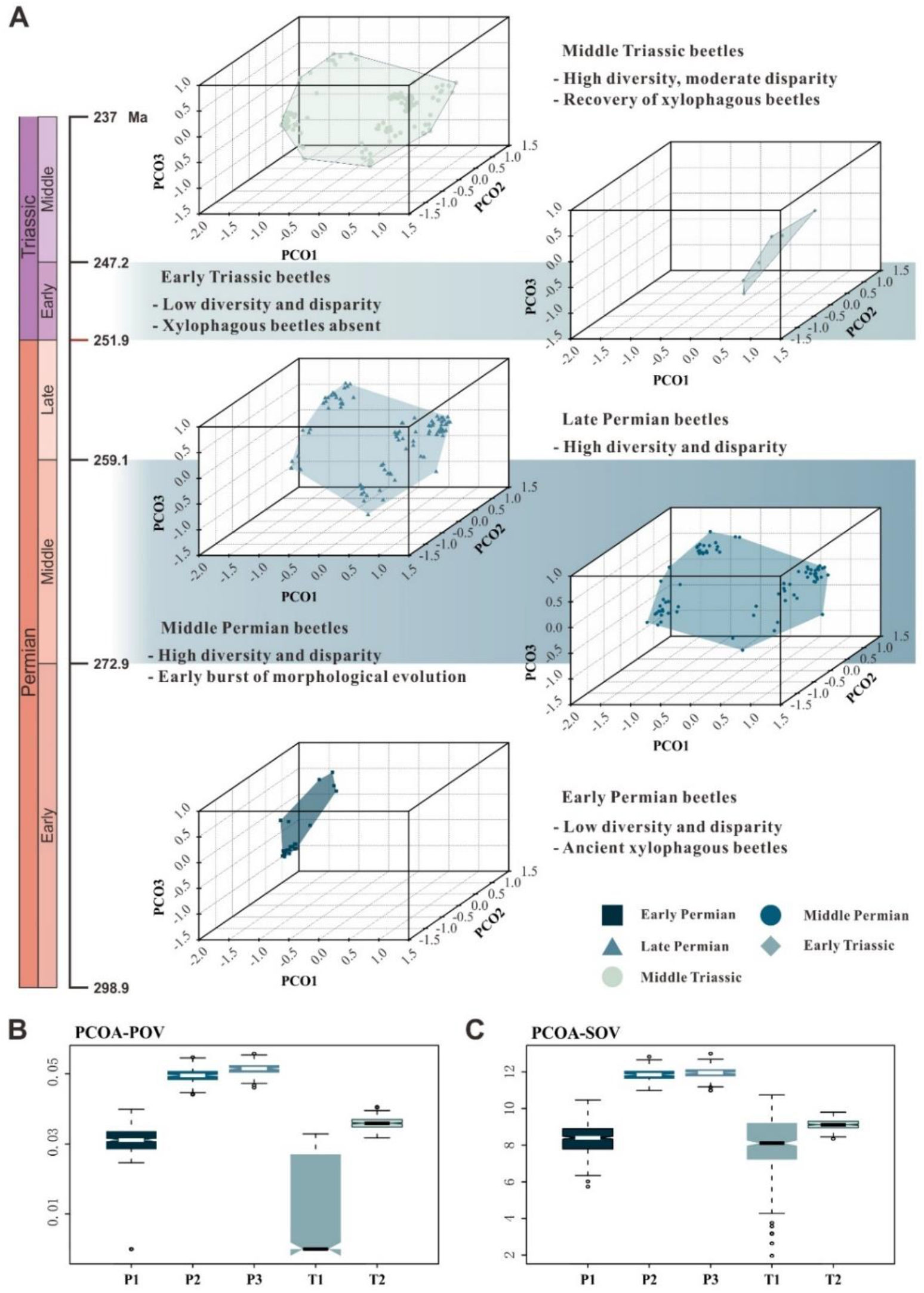
Morphospace comparisons of Coleoptera from the Early Permian to Middle Triassic. (**A**) Morphospace 3D-plot ordinated by PcoA, MORD matrices, based on species-level dataset. (**B** and **C**) Disparity comparisons ordinated by PcoA, MORD matrices, based on species-level dataset, proxy by pov and sov. Abbreviations: pov, product of variance; sov, sum of variance; P1, Early Permian; P2, Middle Permian; P3, Late Permian; T1, Early Triassic; T2, Middle Triassic.

## Discussion

Our results demonstrate that beetles display a steady accumulation of taxonomic diversity throughout the Permian (*Figure 2*). The earliest definite beetles are Tshekardocoleidae from the Early Permian of Germany, Czech Republic, USA and Russia (*Figure 1*), although the origin of Coleoptera is dated to the Carboniferous by molecular phylogenetic analysis (Mckenna et al., 2019). The coleopteran diversity radiation in the Middle Permian is consistent with an expansion of morphological disparity, corresponding to the appearance of multiveined, smooth, and striate elytra as well as some other patterns (*Fig 4* and *Appendix 1—figure 3*). The taxonomic diversity and morphological disparity were decoupled during the Late Permian when taxonomic diversity increased but the morphological disparity was almost stable (*Figures 2, 4* and *Appendix 1—figure 3*). The abrupt Middle Permian increase of coleopteran morphological disparity conforms to the early burst model of clade disparity commonly arising early in radiations (*Simões et al., 2020*; Hughes et al., 2013).

The Permian coleopteran assemblage was dominated by stem groups including Tshekardocoleidae, Permocupedidae (Permocupedinae and Taldycupedinae), and Rhombocolediae in terms of richness and abundance. These ancient beetles were undoubtedly xylophagous because they display a prognathous head, a characteristic elytral pattern with window punctures, a cuticular surface with tubercles (or scales), and a plesiomorphic pattern of ventral sclerites, very similar to the extant wood associated archostematans (*Figure 1*) (Kirejtshuk et al., 2014, Ponomarenko, 1969).

Moreover, Permian trace fossils showing wood-boring provide convincing evidence for the xylophagous habit of these ancient beetles (*Naugolnykh and Ponomarenko, 2010*; Feng et al., 2019). Aquatic or semi-aquatic beetles including Phoroschizidae and Ademosynidae, belonging to the suborder Archostemata, first appeared in the Middle Permian and diversified in the Late Permian (Ponomarenko, 2003). The three other suborders of Coleoptera, comprising Polyphaga, Adephaga and Myxophaga, most likely evolved by the Late Permian, but definite fossils are rare at this time.

Permian beetles probably played an important ecological role in forest ecosystems because most Permian beetles were xylophagous insects that consume living and dead woody stems (*Figure 3*). Some Permian xylophagous beetles fed on living wood tissues (*Feng et al., 2017, 2019*) and could reduce tree productivity and sometimes cause extensive tree mortality, resulting in large transfers of carbon from biomass to dead organic matter (Seidl et al., 2018; Fei et al., 2019). The other Permian xylophagous beetles were saproxylic (feeding on dead wood) (Ponomarenko, 2003), and they could also impact terrestrial carbon dynamics by accelerating wood decomposition (Ulyshen, 2018). Saproxylic animals first appeared in the Devonian and are mainly represented by small invertebrates such as oribatid mites until the Permian (Labandeira et al., 1997; Labandeira, 2007). Whereas grazing by micro- and meso-invertebrates (nematodes, collembolans, enchytraeids and oribatid mites) did not significantly affect wood decomposition, consuming by macro-invertebrates (dominated by saproxylic beetles and termites in modern ecosystems) significantly sped up wood decomposition (*Tapanila and Roberts, 2012*). In addition to those that directly facilitated decomposition by consuming wood, Permian saproxylic beetles are likely to have had a variety of indirect effects on decomposition, including creating tunnels that facilitate the movement of fungi into wood (*Naugolnykh and Ponomarenko, 2010*; Feng et al., 2017), and vectoring fungi and other decay organisms on or within their bodies, like their extant counterparts (Ulyshen, 2016). In conclusion, Permian beetles that feed on living and dead wood probably could impact terrestrial carbon dynamics by reducing forests’ carbon sequestration capacity, and by converting live materials to dead organic matter and subsequent decomposition.

Herbivorous insects and amniotes are thought to be the major herbivorous animals during the Permian and thus are considered to be the most important drivers of the Permian change in biogeochemical cycles of carbon (Laakso et al., 2020). However, Permian herbivorous amniotes mainly fed on leaves, stems, roots and rhizomes (*Sues and Reisz, 1998*; Pearson et al., 2013) and could normally digest cellulose by fermentation but could not consume lignin, as in extant herbivorous vertebrates (Pearson et al., 2013). The majority of terrestrial plant biomass is stored in forest woody tissue consisting of decay-resistant lignin (Hibbett et al., 2016; *Bar-on et al., 2018*). In modern forests, the total carbon stock in woody tissue (including living and dead wood) is about approximately 340 Pg carbon, much more than 72 Pg carbon in roots (below ground), 43 Pg carbon in foliage, and 43 Pg carbon in litter (Reich et al., 2014; Pan et al., 2011). During the Permian, beetles were probably the dominant consumers of woody tissue, while a few other insect groups may have sometimes fed on dead wood (such as stem dictyopterans and protelytropterans) (*Grimaldi and Engel, 2005*). The Early Permian onset of the decrease in oxygen concentrations is consistent with the origin and radiation of the xylophagous beetles in the fossil record. Therefore, we propose that Permian xylophagous beetles could have been responsible for the change of biogeochemical cycles of carbon in Laakso’s model (Laakso et al., 2020).

As the most taxonomically and functionally diverse group of living organisms on Earth (Stork, 2018), extant insects have significant effects on terrestrial carbon and nutrient cycling by modulating the quality and quantity of resources that enter the detrital food web (*Belovsky and Slade, 2000*; *Yang and Gratton, 2014*). However, the effects of insects on terrestrial ecosystems in deep time have been viewed as unimportant or overlooked (Doughty, 2017). Permian beetles were the principal degraders of wood and played a fundamental role in deep-time carbon and nutrient cycling and niche creation. Insects may have been one of the major regulating factors of forest ecosystems at least from the Permian.

Our results show that both the taxonomic diversity and morphological disparity dropped dramatically during the Early Triassic (*Figures 2, 4* and *Appendix 1—figure 3*). Combined with the phylogenetic results (*Figure 3*), our suite of analyses yields a clear ecological signal from beetles across the Permian/Triassic boundary: all xylophagous stem-group beetles become extinct near the Permian-Triassic boundary or abruptly decreased in the Early Triassic [a pattern called “dead clade walking” (Barnes et al., 2021)], while aquatic phoroschizid and ademosynid lineages crossed the Permian/Triassic boundary and diversified in the Middle Triassic. Coleoptera recovered in taxonomic diversity during the Middle Triassic by the rise of new predatory and herbivorous groups, synchronized with the recovery of terrestrial ecosystems (Zhao et al., 2020). However, the morphological disparity is significantly lower than that of the Middle and Late Permian due to the lack of stem-group beetles that possess complex elytra structures (*Figure 4* and *Appendix 1—figure 3*).

Polyphagan groups increased in taxonomic diversity during the Middle Triassic, which is a transitory epoch from a Palaeozoic stem-group beetle assemblage to a Mesozoic polyphagan-dominated assemblage.

Xylophagous groups are absent or rare in Early Triassic coleopteran assemblages, becoming widespread again from the Middle Triassic, mainly represented by more derived archostematans (such as Cupedidae) and polyphagans (Ponomarenko, 2003). This gap in xylophagous beetles coincided chronologically with the gap in coal deposition (“coal gap”) which corresponds to the absence of peat-forming forests (*Figure 3B*) (*Benton and Newell, 2014*, Retallack et al., 1996, Nowak et al., 2020). During the latest Permian and earliest Triassic, gymnosperm-dominated forests abruptly collapsed (Vajda et al., 2020) were replaced by other biomes [such as isoetalean-dominated herbaceous heathlands (Feng et al., 2020)] in most areas due to extreme conditions including aridity (Sun et al., 2012), wildfires (Shen et al., 2011), and ozone destruction (Benca et al., 2018). In some regions the plant extinction was less severe, or the recovery was rapid (Hochuli et al., 2010), or there may have been multiple crises during the Early Triassic (*Schneebeli-Hermann et al., 2017*), but even short-term ecosystem disruption could have led to extinctions among xylophagous beetles. The demise of most forests [deforestation event (Vajda et al., 2020)] most likely resulted in the extinction of most Palaeozoic xylophagous beetles, analogous to the extinction of tree-dwelling birds resulting from end-Cretaceous deforestation (Field et al., 2018).

Our results reveal an Early Triassic gap in xylophagous beetles, suggesting that early archaic beetles experienced the severe ecological consequences of end-Permian deforestation. Extant insects are suffering from dramatic declines in abundance and diversity largely due to the anthropogenic deforestation and global warming (*Van Klink et al., 2020*; Wagner et al., 2021). However, xylophagous insects have been largely neglected in studies of the current extinction crisis (*Van Klink et al., 2020*).

Our findings may help to better understand future changes in insect diversity and abundance faced with global environmental change.

## MATERIALS AND METHODS

### Diversity analysis

We compiled an updated database of all coleopteran species from the Early Permian to Middle Triassic from published literature in the early 1800s through to early 2020. In addition, we incorporated data from other open access database projects, including the Fossil Insect Database (EDNA) and Paleobiology Database (PBDB). We re-examined all published occurrences and taxonomy of Coleoptera from the Early Permian to Middle Triassic (*Supplementary Dataset 1*). We standardized and corrected for nomenclatural consistency of all taxa using a classification of extinct beetle taxa above the genus rank (Bouchard et al., 2011). The data were filtered and cleaned by removing or reassigning illegitimate, questionable, and synonymous taxa and converting local to global chronostratigraphic units (see *Appendix, Supplemental Text D*).

We allocated fossil species into 12 stage-level time bins covering the Early Permian–Middle Triassic interval (from the Asselian to Ladinian, 298–237 Ma). Considering the short duration of the Induan and Olenekian stages, we combined both stages into one time bin. The formal taxa were erected based only on isolated elytra that cannot be classified definitely into any natural group. Thus, we separately counted the diversity of natural, formal and mixed groups (*Figure 2* and *Appendix 1— figure 1*). We determined the stratigraphical ranges of families, genera and species as the maximum and minimum ages in stage-level time bins. All diversity was calculated using the range-through method.

### Phylogenetic analysis

In light of the new taxa and characters available for further testing the phylogenetic status of ancient stem-group beetles, we reconstructed the phylogenetic relationships among the stem groups by incorporating the presently described new taxa and revised characters coding into the previous dataset (Beutel et al., 2008). The morphological characters used for phylogenetic analysis comprise 93 adult and larval characters (*Supplementary Dataset S2*). Unknown characters were coded as ‘?’. The taxon sampling contains two megalopterans as outgroups (*Sialis* and *Chauliodes*) and 13 coleopteran ingroup taxa (five extant and 8 extinct) representing all four coleopteran extant suborders and their stem groups. Compared to previous character matrices, we added the subfamily Taldycupedinae and three new characters. The matrix was analysed in TNT ver. 1.1, through parsimony analysis and using traditional search (Goloboff et al., 2008). All characters were equally weighed and unordered (1000 replicates and 1000 trees saved per replication). Bootstrap values, consistency index (CI) and retention index (RI) were provided (*Appendix 1—figure 2*).

### Morphospace analysis

We performed morphospace analyses with our newly assembled discrete character matrices (*Supplementary Dataset S3*). The analyses were performed using the free software R.4.0.4. Both the Maximum Observable Rescaled Distance (MORD) matrix and Generalized Euclidean Distance (GED) matrix were calculated based on two discrete character matrices (Lloyd, 2016; Wills, 1998). Recent research has revealed that the GED matrix creates a systematic bias in cases with a high percentage of missing data, and the MORD matrix can provide a greater fidelity under these circumstances (Lloyd, 2016; Lehmann et al., 2019). We then ordinated all taxa into a multivariate morphospace with both principal coordinate analysis (PcoA) and non-metric multidimensional scaling (NMDS).

For PcoA, we used the function “ordinate cladistic matrix” in package “Claddis” with a cailliez method to correct the negative eigenvalues (Lloyd, 2016). Two disparity matrices were used to evaluate the volume of the morphospace, including the sum and product of the variances (Wills et al., 1994). The product metrics was normalized by taking the nth root (n equals the number of axes used for calculating disparity metrics). We used the scores on all axes that together comprise 90% of total variance to calculate those disparity metrics. We chose a permutation test (two-tailed) to test the null hypothesis of no difference between insect disparity of different time-bins. Each test run used 5000 replications. The test statistic was obtained by using the disparity metric of an older time-bin to minus that of a younger time-bin. If the proportion in the null distribution greater than the observed value of the test statistic is smaller than 0.025, the insect disparity of an older time-bin was considered significantly larger than that of a younger time-bin, and if the proportion was greater than 0.975, the insect disparity of a younger time-bin was considered significantly larger than that of an older time-bin (*Supplementary Dataset S4*).

We also performed permutation tests with sample size corrected. For two design groups with different sample sizes, we first performed subsampling of the group with more samples to obtain equal sample sizes. Based on the newly obtained two groups with equal sample sizes, we calculated the observed value of the test statistic. Then we randomly permutated those species into different groups once and calculated a test statistic. We repeated this procedure (subsampling and permutation) 10000 times and obtained a null distribution plus a group of observed values. We then calculated a set of proportions greater than the observed values in the null distribution. By analogy, if the median of the proportions is equal to or smaller than 0.025, the insect disparity of an older time-bin is significantly larger than that of a younger time-bin. If the median proportion is considered significantly greater than 0.975, the insect disparity of a younger time-bin is larger than that of an older time-bin (*Appendix 1—figures 6*–*9*).

For NMDS, we used the function “metaMDS” in package “vegan” with the number of dimension settings to 3 (Dixon, 2003). Both non-metric fit and linear fit were very high (larger than 0.90; *Appendix 1—figures 6–9*) and the stresses were smaller than 0.2 which implies that the ordinations are relatively good. Then we repeated all the previous analyses with this NMDS morphospace and acquired a similar result. The two disparity-metrics of each time-bin were calculated based on two different distance matrices and two different ordination methods (*Figure 4* and *Appendix 1—figures 3–7*). The distribution was simulated under 500 bootstraps.

Thirty-one undetermined specimens from the Grès à Voltzia Formation (Lower/Middle Triassic boundary, France) were included in our database and their age was attributed to the early Middle Triassic in our analysis (*Figure 4* and *Appendix—figures 3–7*). Considering that the age of these specimens is controversial, we repeated all our analyses assuming that the age of these specimens is Early Triassic; the result is consistent with the previous one (*Appendix 1—figures 8–9*).

## Acknowledgements

We thank A.G. Ponomarenko, A.P. Rasnitsyn, A.G. Kirejtshuk, J. Xue, H. Xu, B. Huang, and H. Zeng for helpful discussion, and D. Yang for reconstructions. BW thanks members of the palaeoentomological laboratory of the Palaeontological Institute (Russian Academy of Sciences) for their help during his visit to Moscow (2010).

## Author contributions

B.W. designed the project. Xianye Zhao, Y.Y., and B.W. wrote the manuscript. Xianye Zhao, Y.Y., E.Y., and B.W. prepared the figures. Xianye Zhao, Y.Y., M.E.C., and B.W. performed the comparative and analytical work. Xianye Zhao and Y.Y. performed the morphospace analyses. Xianye Zhao, J.C., and B.W. ran the phylogenetic analyses. Xianye Zhao, E.Y., E.A.J., and Xiangdong Zhao collected data and contributed to the discussion. All authors discussed and approved the final manuscript.

## Data and materials availability

All data needed to evaluate the conclusions in the paper are present in the paper and/or the Supplementary Materials. Additional data related to this paper may be requested from the authors.

## Supplementary Materials

### Appendix Supplementary Text

#### A. List of taxa used for the phylogenetic analysis

Order Megaloptera

Family Sialidae Leach, 1815

1 Genus *Sialis* Latreille, 1803

Family Chauliodidae Davis, 1903

2 Genus *Chauliodes* Latreille, 1796

Order Coleoptera

Suborder Polyphaga Emery, 1886

Family Hydrophilidae Latreille, 1802

3 Genus *Helophorus* Fabricius, 1775

Suborder Myxophaga Crowson, 1955

Family Torridincolidae Steffan, 1964

4 Genus *Torridincola* Steffan, 1964

Suborder Adephaga Clairville, 1806

Family Trachypachidae Thomson, 1857

5 Genus *Trachypachus* Motschulsky, 1845

Suborder Archostemata Kolbe, 1908

Family Cupedidae Laporte, 1836

6 Genus *Cupes* Fabricius, 1801

Family Ommatidae Sharp and Muir, 1912

7 Genus *Omma* Newman, 1839

Family Tshekardocoleidae Rohdendorf, 1944

8 Genus *Tshekardocoleus* Rohdendorf, 1944

Family Permocupedidae Martynov, 1932

Subfamily Permocupedinae Martynov, 1932

9 Genus *Permocupes* Martynov, 1932

Subfamily Taldycupedinae Rohdendorf, 1961

10 Genus *Taldycupes* Rohdendorf, 1961

Family Rhombocoleidae Rohdendorf, 1961

11 Genus *Rhombocoleites* Ponomarenko, 1969

Family Triadocupedidae Ponomarenko, 1966

12 Genus *Triadocupes* Ponomarenko, 1966

Family Ademosynidae Ponomarenko, 1968

13 Genus *Chaocoleus* Ponomarenko, Yan & Huang, 2014

Family Phoroschizidae Bouchard and Bousquet, 2020

14 Genus *Dikerocoleus* Lin, 1982

Family Triaplidae Ponomarenko, 1977

15 Genus *Triaplus* Ponomarenko, 1977

#### B. Characters used for the phylogenetic analysis

1. Externally visible membranes: (0) present; (1) absent.
2. Tubercles: (0) absent or very indistinct; (1) present.
3. Scale-like setae: (0) absent; (1) present.
4. Ocelli: (0) three; (1) absent.
5. Constricted neck and postocular extensions: (0) absent or indistinct; (1) present.
6. Supraantennal protuberance: (0) absent; (1) present as moderately distinct bulge;
7. (2) present as strongly pronounced protuberance.
8. Supraocular protuberance: (0) absent; (1) present as moderately distinct bulge; (2) present as strongly pronounced protuberance.
9. Posteromesal protuberance: (0) absent; (1) present, moderately convex; (2) conspicuous, strongly convex.
10. Posterolateral protuberance: (0) absent; (1) present.
11. Antennal groove on head; (0) absent; (1) below compound eye; (2) above compound eye.
12. Gular sutures: (0) complete, reaching hind margin of head capsule; (1) incomplete, not reaching hind margin of head capsule; (2) absent.
13. Shape of gula: (0) not converging posteriorly; (1) converging posteriorly.
14. Tentorial bridge: (0) present; (1) absent.
15. Posterior tentorial grooves: (0) externally visible; (1) not visible externally.
16. Anterior tentorial arms: (0) well developed; (1) strongly reduced or absent, not connected with posterior tentorium.
17. Frontoclypeal suture: (0) present; (1) absent.
18. Labrum: (0) free, connected with clypeus by membrane; (1) indistinctly separated from clypeus, largely or completely immobilized; (2) fused with head capsule.
19. Musculus labroepipharyngalis: (0) present; (1) absent.
20. . Musculus frontolabralis: (0) present; (1) absent.
21. Musculus frontoepipharyngalis: (0) present; (1) absent.
22. Length of antenna: (0) not reaching mesothorax posteriorly; (1) very elongate, reaching middle region of body.
23. Number of antennomeres: (0) 13 or more; (1) 11 or less.
24. Location of antennal insertion on head capsule: (0) laterally; (1) dorsally.
25. Extrinsic antennal muscles: (0) four; (1) three; (2) two.
26. Shape of mandible: (0) short or moderately long, largely covered by labrum in repose (1) very elongate and protruding in resting position (3) vestigial.
27. Ventromesal margin of sculptured mandibular surface: (0) not reaching position of mandibular condyle; (1) reaching mandibular condyle.
28. Cutting edge of mandible: (0) horizontal (1) with three vertically arranged teeth.
29. Separate areas with different surfaces on ventral side of mandible; (0) absent; (1) present.
30. Deep pit in cranio-lateral area of ventral surface of mandible: (0) absent; (1) present.
31. Galea: (0) without globular distal galeomere and basal galeomere not slender and stalk-like; (1) stalk-like basal galeomere and globular distal galeomere; (2) absent.
32. Lacinia: (0) present; (1) absent.
33. Apical segment of maxillary palp: (0) with only one apical field of sensilla (campaniform sensilla) (1) with an apical and a dorsolateral field of sensilla.
34. Digitiform sensilla on apical maxillary palpomere: (0) absent; (1) present.
35. Pit containing sensilla on dorsolateral field of apical maxillary palpomere: (0) absent; (1) present.
36. Deep basal cavity of prementum: (0) absent; (1) present.
37. Lid-like ventral premental plate: (0) absent; (1) present.
38. Transverse ridge of prementum: (0) absent; (1) present.
39. Anterior appendages of prementum: (0) paired ligula; (1) ligula subdivided into many digitiform appendages; (2) absent.
40. Mentum: (0) distinctly developed; (1) vestigial but recognizable as a transverse sclerite between the submentum and the premental plate; (2) absent.
41. Musculus tentoriopharyngalis posterior: (0) moderately sized, not distinctly subdivided into individual bundles; (1) complex, composed of series of bundles, origin from the gular ridges or lateral gular region.
42. Propleural suture (0) present; (1) absent.
43. Exposure of propleura: (0) fully exposed, propleura reaches anterior margin of prothorax; (1) exposed, not reaching anterior margin of prothorax; (2) internalized.
44. Fusion of propleura and protrochantinus: (0) absent; (1) present.
45. Prosternal grooves for tarsomeres: (0) absent; (1) present.
46. Length of prosternal process: (0) not reaching beyond hind margin of procoxae, very short or absent; (1) reaching hind margin of procoxae.
47. Shape of prosternal process: (0) not broadened apically; (1) apically broadened and truncate.
48. Broad prothoracic postcoxal bridge: (0) absent; (1) present.
49. Mesocoxal cavities: (0) not bordered by metanepisterum; (1) bordered by metanepisternum.
50. Mesoventrite with anteromedian pit for reception of prosternal process: (0) absent or only very shallow concavity; (1) distinct, rounded groove; (2) large hexagonal groove.
51. Propleuro-mesepisternal locking mechanism: (0) absent; (1) propleural condyle and mesepisternal socket; (2) mesepisternal condyle and propleural socket.
52. Connection of meso- and metaventrite: (0) sclerites distinctly separated, connected by a membrane; (1) articulated but not firmly connected; (2) firmly connected between and within mesocoxal cavities.
53. Transverse suture of mesoventrite: (0) present; (1) absent.
54. Mesal coxal joints of mesoventrite: (0) present; (1) absent.
55. Shape of mesocoxae: (0) globular or conical; (1) with deep lateral excavation and triangular lateral extension.
56. Exposed metatrochantin: (0) present, distinctly developed; (1) indistinct or absent.
57. Shape of penultimate tarsomere: (0) not distinctly bilobed; (1) distinctly bilobed.
58. Forewings: (0) membranous; (1) transformed into sclerotized elytra.
59. Venation of forewings: (0) distinct, not arranged in parallel rows; (1) parallel arrangement of distinct longitudinal veins; (2) longitudinal veins very indistinct or absent.
60. Elytral sclerotization pattern: (0) with a pattern of unsclerotized window punctures; (1) entirely sclerotized.
61. Elytral apex: (0) distinctly reaching beyond abdominal apex posteriorly; (1) slightly reaching beyond abdominal apex posteriorly; (2) reaching abdominal apex or shorter.
62. Transverse folding mechanism of hind wings: (0) absent; (1) present.
63. Oblongum cell of hind wing: (0) closed cell not differentiated as oblongum cell;
64. oblongum present; (2) open or absent.
65. Abdominal sternite I: (0) exposed; (1) concealed under metacoxae, largely or completely reduced.
66. Median ridge on ventrite 1: (0) absent; (1) present.
67. Number of exposed abdominal sternites (excluding sternite I): (0) more than six;
68. six; (2) five.
69. Arrangement of abdominal sterna: (0) abutting, not overlapping; (1) tegular or overlapping.
70. Difference between main vein and interval vein: (0) significant; (1) unconspicuous.
71. Number of window punctures rows: (0) more than 11 rows; (2) 11 rows; (3) less or equal to 10 rows.
72. Multiple rows of window punctures in the base of elytron (more than 2 rows): (0) present; (1) absent.
73. Head shape of later instars: (0) parallel-sided, slightly narrowing anteriorly, or evenly rounded; (1) transverse, strongly rounded laterally, greatest width near hind margin.
74. Posteromedian emargination of head capsule: (0) absent; (1) present.
75. Endocarina: (0) absent; (1) present, undivided; (2) present, forked.
76. Frontal suture of second and third instars: (0) distinct; (1) indistinct or absent.
77. Stemmata: (0) more than one pair of stemmata; (1) one pair of stemmata or eyeless.
78. Length of antenna: (0) at least 20% of greatest width of head capsule; (1) less than 20% of greatest width of head capsule.
79. Antennal segments: (0) four or more; (1) three or less.
80. Shape of distal part of mandible: (0) less than three apices; (1) three apices.
81. Retinaculum: (0) present; (1) absent.
82. Shape of mola: (0) not quadrangular, not delimited by a distinct margin; (1) quadrangular and delimited by a distinct margin; (2) missing.
83. Ligula: (0) unsclerotized; (1) sclerotized, enlarged and wedge-shaped.
84. Mentum and submentum: (0) not fused; (1) fused and narrowed between maxillary grooves.
85. Prothorax: (0) as broad as following segments; (1) broader than following segments.
86. Leg segments: (0) six; (1) five.
87. Leg segments: (0) six; (1) five.
88. Abdominal segments I–III of later instars: (0) shorter than thorax; (1) longer than thorax.
89. Tergal ampullae: (0) absent; (1) present.
90. Ventral asperities: (0) absent; (1) present.
91. Lateral longitudinal bulge of abdominal segments I–VII: (0) absent; (1) present.
92. Sclerotized process of tergum IX: (0) absent; (1) present.
93. Eversible lobes of segment IX: (0) absent; (1) present.
94. Urogomphi: (0) absent; (1) present.
95. Segment X: (0) exposed; (1) not visible externally.
96. Larval habitat: (0) not associated with wood; (1) associated with wood.

#### C. Characters used for the morphospace analysis

1. Sides of elytron: (0) external margin more convex;(1) sutural margin more convex;
2. both similar.
3. Shape of window punctures: (0) polygonal; (1) round or oval; (2) without window punctures.
4. M vein and Cu vein: (0) forming X-shaped vein; (1) not forming X-shaped vein; (2) without veins.
5. Number of impression row(s): (0) without impression row; (1) 7 rows; (2) 8 rows;
6. 9 rows.
7. Striae of elytron: (0) striae with punctures; (1) striae without punctures; (2) elytron without striae.
8. Schiza present or absent: (0) present; (1) absent.
9. Number of puncture rows: (0) elytron without puncture row; (1) 3 rows; (2) 7 or 8 rows; (3) 12 rows.
10. Shape of elytron: (0) widest at base part of elytron; (1) middle of elytron widest; (2) the base and middle similarly wide; (3) widest at apex of elytron.
11. Short striae of the base of elytron: (0) present; (1) absent.
12. Fusion of striae in the termination of elytron: (0) fusion of striae present; (1) striae not fused at apex; (2) elytron without striae.
13. Striae elongated to the termination: (0) present; (1) absent; (2) elytron without striae.
14. Lateral location of schiza: (0) on the base of elytron; (1) on the middle of elyton;
15. on the apex of elytron; (3) without absent.
16. Denticles of elytral margin: (0) present; (1) absent.
17. Longitudinal location of schiza: (0) near the external margin; (1) on the middle of elytron; (2) near the sutural margin; (3) schiza absent.
18. Interval vein: (0) mainly zigzag vein; (1) mainly straight vein; (2) mixed with zigzag and straight veins; (3) without veins.
19. Difference between main vein and interval vein: (0): significant; (1) blurry; (2) without vein.
20. Elytral veins elongated to the termination (excluding C, SC, A vein): (0) present;
21. absent; (2) elytron without veins.
22. Elytron tail: (0) present; (1) absent.
23. Branches of vein: (0) present; (1) absent; (2) elytron without branches.
24. Number of tubercles rows: (0) tubercle rows absent; (1) less than or equal to 7 rows; (2) more than 7 rows.
25. Tubercles rows between ridges: (0) present; (1) tubercle rows present but without ridges; (2) elytron without tubercle row.
26. Elytra aspect ratio: (0) less than 3; (1) 3-4; (2) more than 4.
27. Tubercles: (0) present; (1) absent.
28. Discrete punctures (or pits): (0) present; (1) absent.
29. Epipleuron: (0) absent; (1) present.
30. Epipleural border: (0) absent; (1) present.
31. Sutural border: (0) present; (1) absent.
32. Number of puncture rows in epipleuron: (0) single row; (1) multiple rows; (2) epipleuron without puncture rows; (3) elytron without epipleuron.
33. Punctures (or pits) in epipleural border: (0) present; (1) absent; (2) elytron without epipleural border.
34. Punctures (or pits) in sutural border: (0) present; (1) absent; (2) elytron without epipleural border.
35. Number of striae: (0) less or equal to 10 rows; (1) more than 10 rows.
36. Ribs: (0) present; (1) absent.
37. Multiple rows of window punctures in base of elytron (more than 2 rows): (0) present; (1) absent; (2) elytron without window punctures.
38. Adjacent multiple short striae in base of elytron: (0) present; (1) absent; (2) elytron without striae.
39. Shallow striae: (0) present; (1) absent.

#### D. Taxonomic revisions

Revisions for Taldycupedinae.

The subfamily Taldycupedinae 1961 was characterized by the absence of branching vein, 7 to 10 parallel windows punctures, main vein similar to the intermediate veins and some abdominal characters (Wagner et al., 2021). Later, Ponomarenko revised the diagnosis of the family as ten window puncture rows, with additional cell rows basally in most cases and 3 window puncture rows behind A3 (Ponomarenko, 1969). And then Carpenter removed the character “3 window puncture rows behind A3” from the diagnosis (Carpenter, 1992). Moreover, some species reported by Ponomarenko are not consistent with the diagnosis of ten window puncture rows, such as *Tecticupes martynovi* and *Tychticupoides grjasevi* (Ponomarenko, 1969; Aristov et al., 2013). Thus, the number of window puncture rows is not a definite character for this family. The basal veins, especially those in the cubital area, are unique among stem groups (Tshekardocoleidae, Permocupedinae, and Taldycupedinae) as an important character which is absent in extant beetles. Thus, we consider that some taxa of Taldycupedinae without this character should be re-examined. Here, we re-examined Mesozoic Taldycupedinae and proposed that the following specimens should be removed from this subfamily.

1.*Mesothoris grandis* Dunstan, 1923

This species was reported from the Upper Triassic of Queensland, Australia (Dunstan, 1923). The base of the holotype is lacking. Thus, we cannot identify whether there are additional windows puncture rows basally. We consider this species cannot be definitely classified to Taldycupedinae.

2.*Mesothoris punctomarginum* Dunstan, 1923

This species was reported from the Upper Triassic of Queensland, Australia (Dunstan, 1923; Kirejtshuk, 2020). We found that the decorations of the elytron are not windows punctures but are similar to rounded tubercles (Jell, 2004). In addition, due to the lack of the base of the elytron, it is not clear whether there are additional cell rows basally, an important character for Taldycupedinae (Ponomarenko, 1969; Carpenter, 1992). Thus, we suggest that this species cannot be definitely classified in Taldycupedinae.

3.*Simmondsia cylindrica* Dunstan, 1923

This species was reported from the Upper Triassic of Queensland, Australia (Dunstan, 1923). We found that there are not distinct additional cell rows basally (Jell, 2004). Besides, there are less than three cell rows in the anal area (Jell, 2004). Thus, we consider this species cannot be definitely classified in Taldycupedinae.

4.*Simmondsia subpiriformis* Dunstan, 1923

This species was reported from the Upper Triassic of Queensland, Australia (Dunstan, 1923). We found that there are not distinct additional cell rows basally (Jell, 2004). In addition, the vein A3 is extended to the elytra apex, that is quite different from the short A3 in other Taldycupedinae (Ponomarenko, 1969; Carpenter, 1992). Thus, we consider this species cannot be definitely classified in Taldycupedinae.

5.*Wuchangia latilimbata* Hong, 1985

This species was reported from the Lower Jurassic of Hubei, China (Hong, 1985a).

The base of the elytron is blurred, and it is not clear whether additional cell rows existed. Besides, this specimen lacks the short anal vein. Thus, we suggest that this species cannot be classified in Taldycupedinae.

6.*Yuxianocoleus hebeiense* Hong, 1985

This species was reported from the Lower Jurassic of Hubei, China (Hong, 1985b).

Its cell punctures are small, similar to striae (small punctures in a thin furrow), not real windows punctures. Thus, we consider that this species cannot be classified as Taldycupedinae.

Revisions for Asiocoleidae.

The family Asiocoleidae contains 12 genera and 28 species from the Middle Permian to Late Triassic of Russia, China, Australia, Kazakhstan, Kyrgyzstan and Mongolia (Ponomarenko, 2011). Ponomarenko (2011) proposed that the family Tricoleidae should be considered a junior synonym of Asiocoleidae (Ponomarenko, 2011). However, we think it is incorrect to merge Tricoleidae with Asiocoleidae because there is no definite autapomorphy for combining these two families.

Therefore, we retain these two families in our analysis. We attributed *Schizotaldycupes*, *Asiocoleus*, *Asiocoleopsis*, *Bicoleus*, and *Tetracoleus* in Asiocoleidae, a formal group that lacks detailed body characters.

Replacement name for *Uskatocoleus convexus* Ponomarenko, 2013.

We noticed that the species name *Uskatocoleus convexus* Ponomarenko, 2013 had already been used by Rohdendorf (*Rohdendorf, et al., 1961*). Therefore, we proposed a new species name *Uskatocoleus ponomarenkoi* as a replacement name for the preoccupied and junior homonym *Uskatocoleus convexus* Ponomarenko, 2013. The new species name is named in honour of the palaeoentomologist A.G. Ponomarenko for his comprehensive and profound contribution to the taxonomy of fossil beetles.

**Appendix figure 1.**
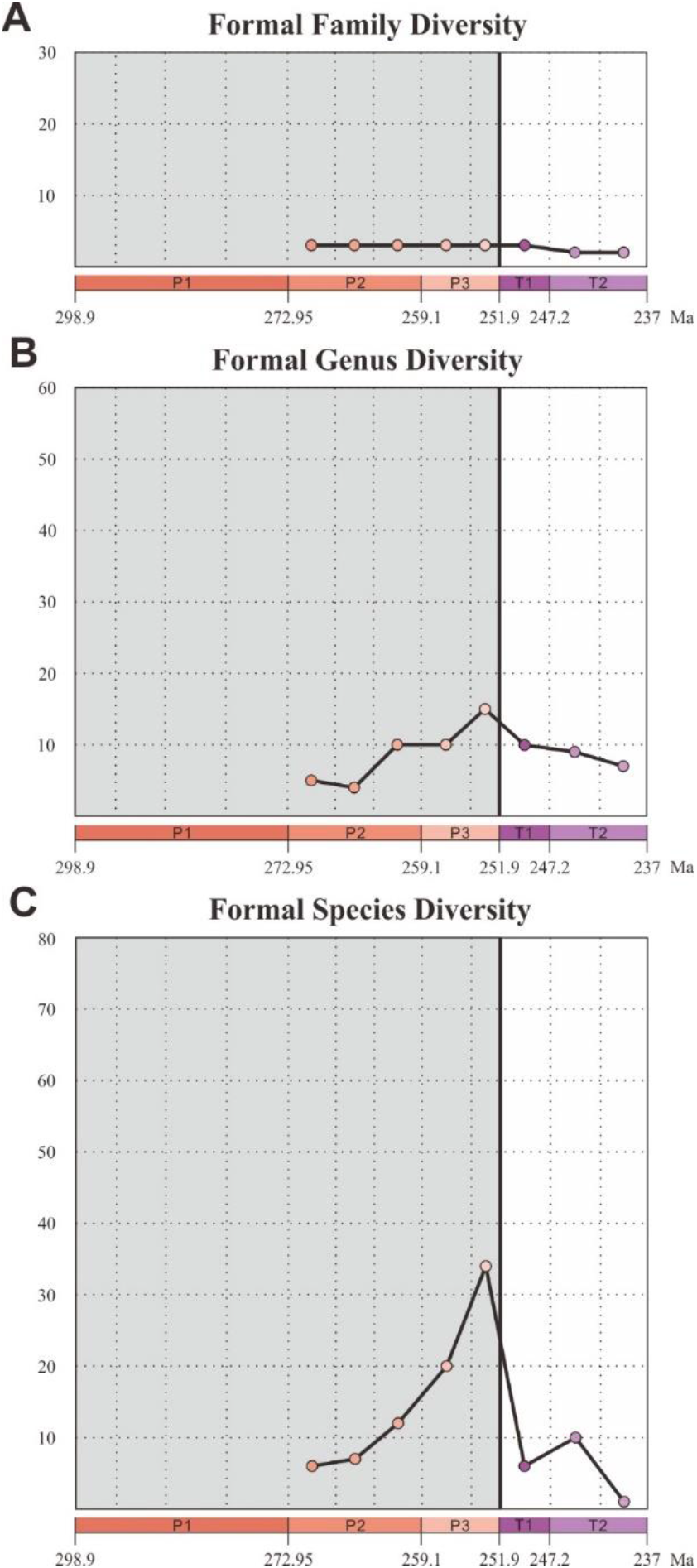
Diversity of Coleoptera formal groups from the Early Permian to Middle Triassic. (**A**) Family-level diversity. (**B**) Genus-level diversity. (**C**) Species-level diversity. Abbreviations: P1, Early Permian; P2, Middle Permian; P3, Late Permian; T1, Early Triassic; T2, Middle Triassic.

**Appendix figure 2.**
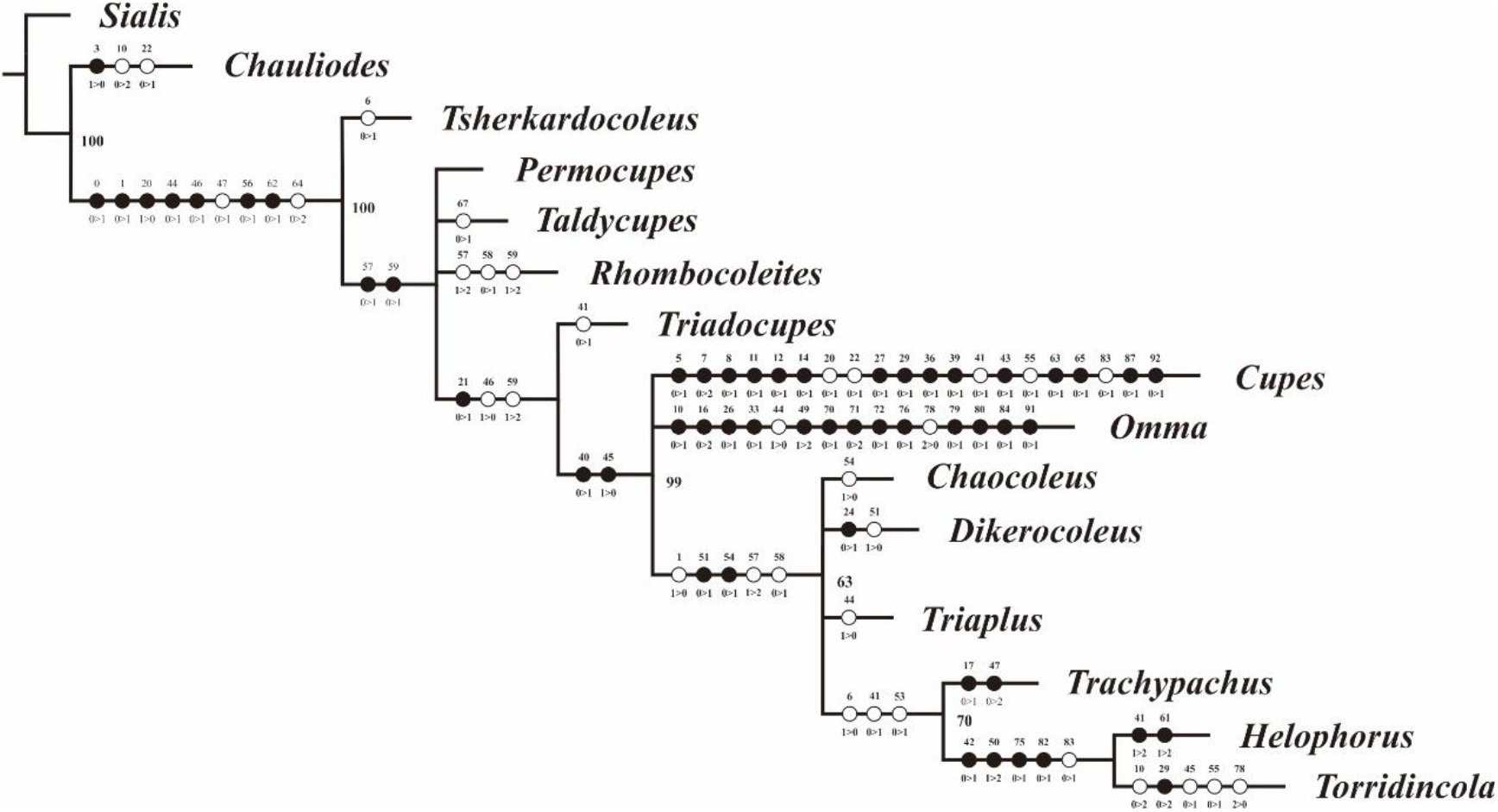
Strict consensus tree of 6 most parsimonious trees of Coleoptera. Tree length = 199, consistency index (CI) = 0.782, retention index (RI) = 0.750. Numbers on branches denote bootstrap frequencies; bootstrap frequencies below 50 are not shown. The unambiguous apomorphies are mapped on the tree (numbers above branches indicate character numbers, and numbers below branches represent state changes).

**Appendix figure 3.**
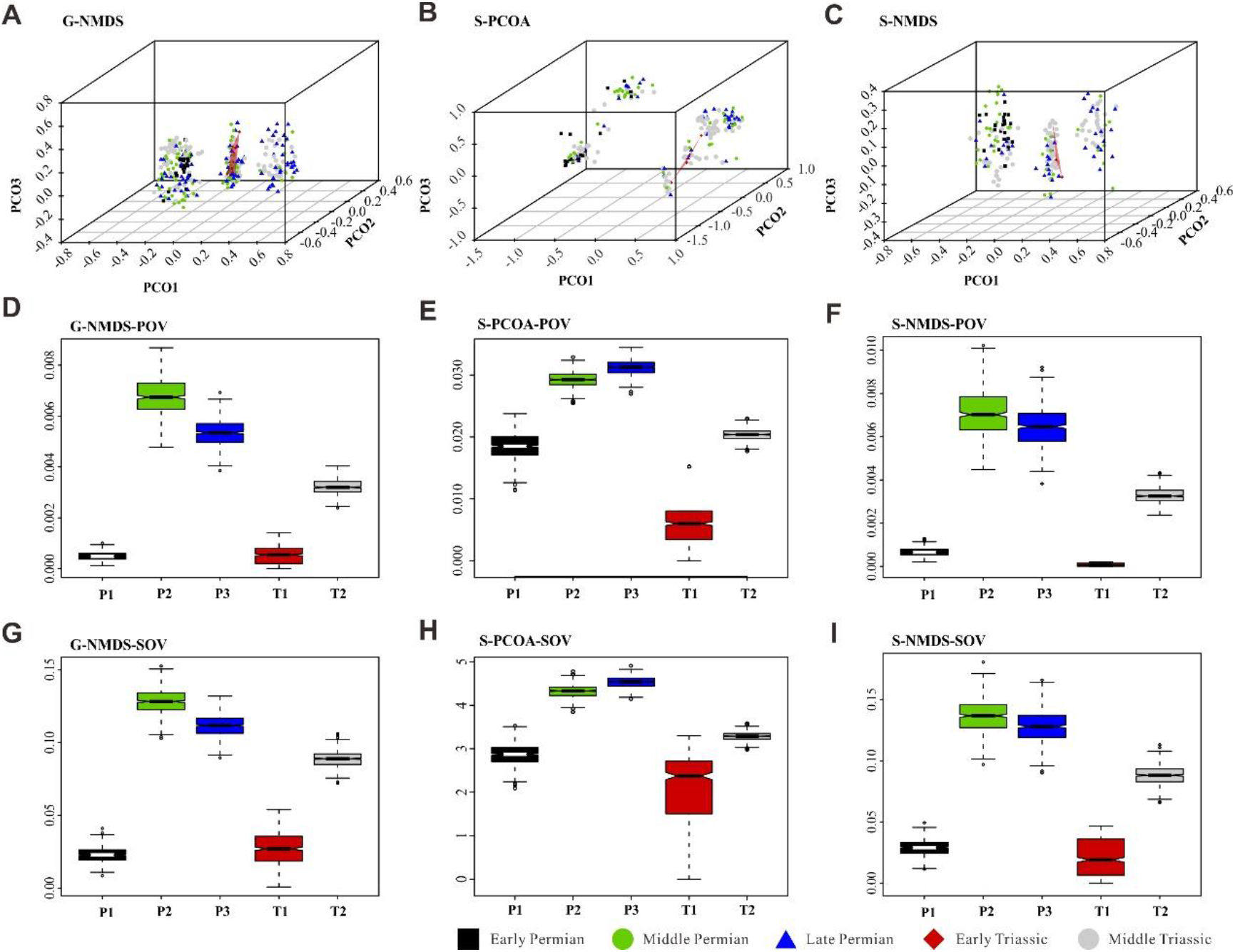
Morphospace comparisons of Coleoptera from the Early Permian to Middle Triassic, MORD matrices, proxy by pov and sov. (A, D and G) Morphospace 3D-plot and disparity comparisons ordinated by NMDS, based on genus-level dataset. (**B**, **E** and **H**) Morphospace 3D-plot and disparity comparisons ordinated by PCOA, based on species-level dataset. (**C**, **F** and **I**) Morphospace 3D-plot and disparity comparisons ordinated by NMDS, based on species-level dataset. Abbreviations: pov, product of variance; sov, sum of variance; P1, Early Permian; P2, Middle Permian; P3, Late Permian; T1, Early Triassic; T2, Middle Triassic. In the morphospace 3D-plot, morphospace occupation of Early Triassic taxa is highlighted by red area.

**Appendix figure 4.**
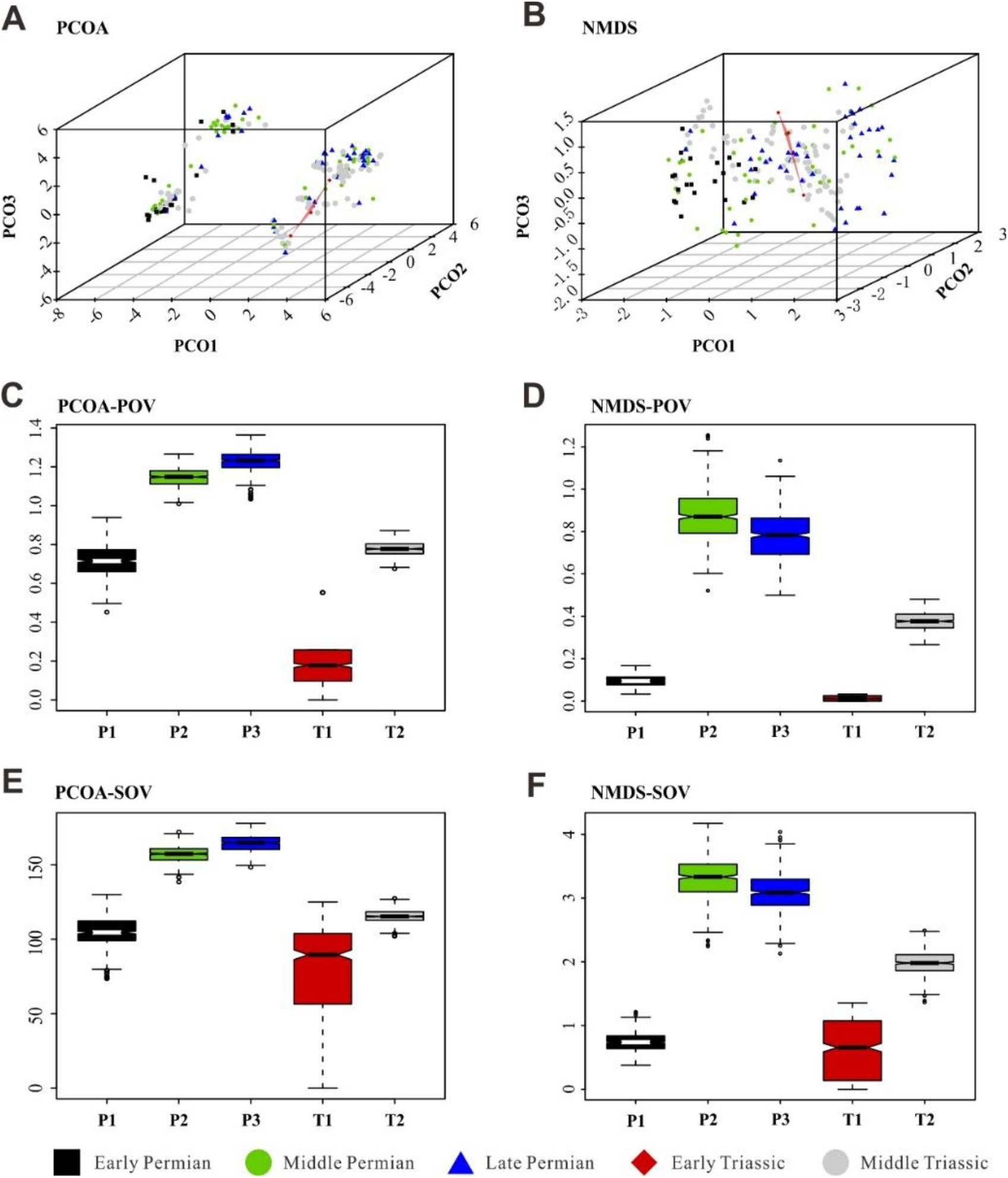
Morphospace comparisons of Coleoptera from the Early Permian to Middle Triassic, based on genus-level disparity analyses of GED matrices, proxy by pov and sov. (**A, C and E**) Morphospace 3D-plot and disparity comparisons ordinated by PCOA. (**B, D and F**) Morphospace 3D-plot and disparity comparisons ordinated by NMDS. Abbreviations: pov, product of variance; sov, sum of variance; P1, Early Permian; P2, Middle Permian; P3, Late Permian; T1, Early Triassic; T2, Middle Triassic. In the morphospace 3D-plot, morphospace occupation of Early Triassic taxa is highlighted by red area.

**Appendix figure 5.**
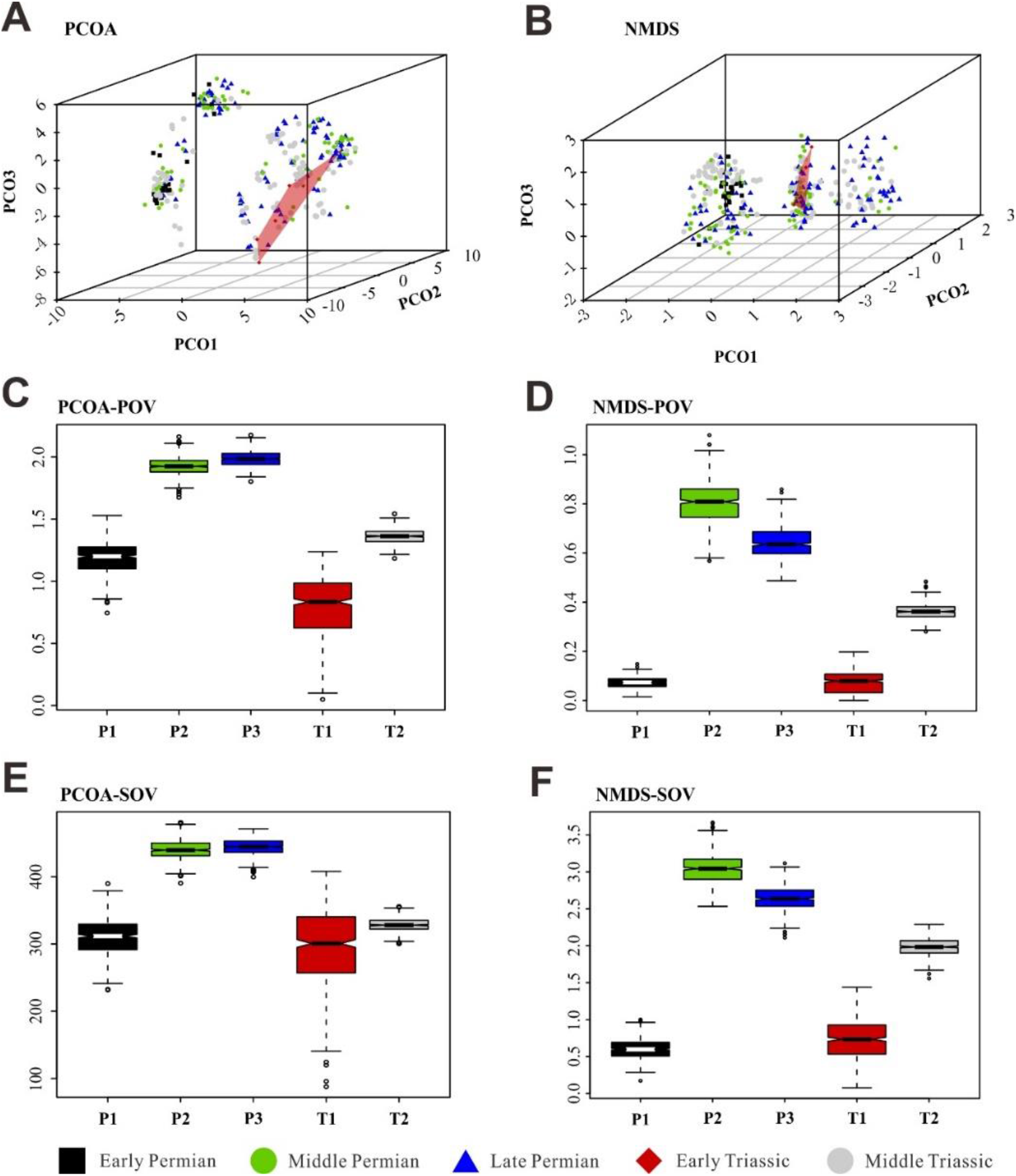
Morphospace comparisons of Coleoptera from the Early Permian to Middle Triassic, based on species-level disparity analyses of GED matrices, proxy by pov and sov. (**A, C and E**) Morphospace 3D-plot and disparity comparisons ordinated by PCOA. (**B, D and F**) Morphospace 3D-plot and disparity comparisons ordinated by NMDS. Abbreviations: pov, product of variance; sov, sum of variance; P1, Early Permian; P2, Middle Permian; P3, Late Permian; T1, Early Triassic; T2, Middle Triassic. In the morphospace 3D-plot, morphospace occupation of Early Triassic taxa is highlighted by red area.

**Appendix figure 6.**
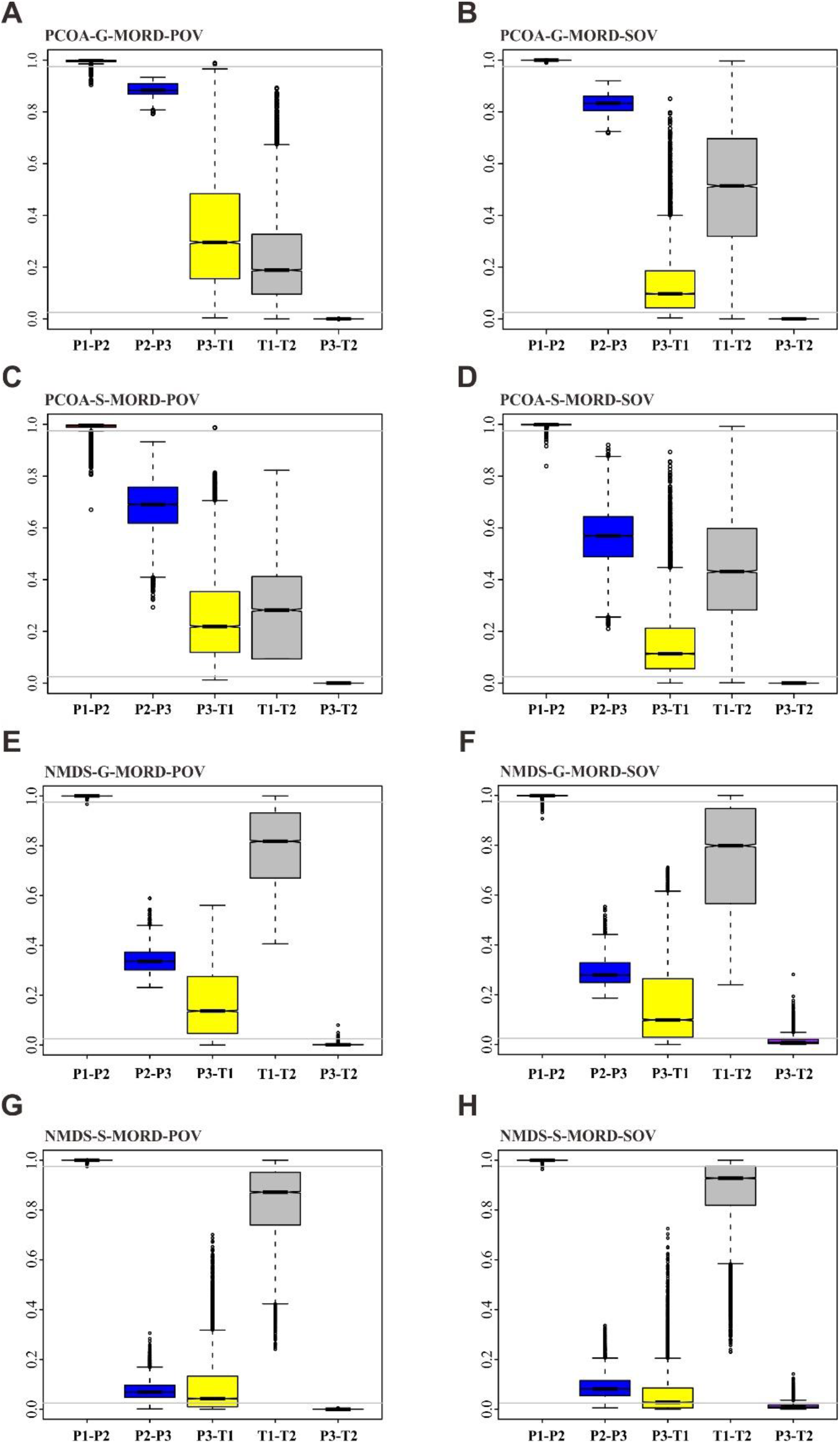
Permutation tests with sample size corrected (MORD). (**A-D**) Permutation tests for PCOA results. (**E-H**) Permutation tests for NMDS results. Each box represents a distribution of the proportion greater than the observed value of the test statistic in the null distribution. Grey line highlights the value of 0.025 or 0.975. Difference lower than 0.025 represents the disparity of the former lower than the latter significantly, and the condition of the difference higher than 0.975 was inverse. Abbreviations: G, genus database; S, species database; pov, product of variance; sov, sum of variance; P1, Early Permian; P2, Middle Permian; P3, Late Permian; T1, Early Triassic; T2, Middle Triassic.

**Appendix figure 7.**
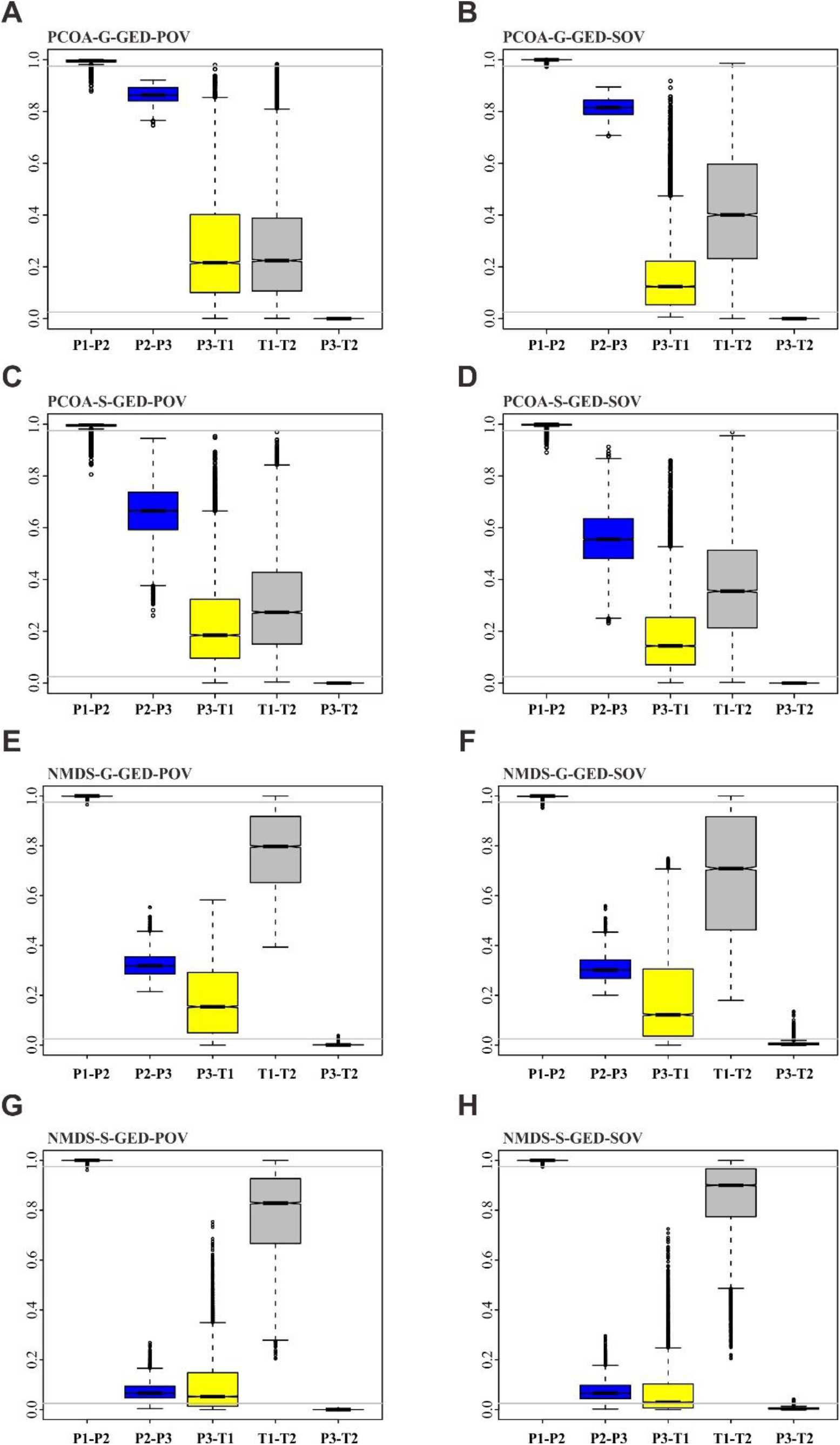
Permutation tests with sample size corrected (GED). (A-D) Permutation tests for PCOA results. (**E-H**) Permutation tests for NMDS results. Each box represents a distribution of the proportion greater than the observed value of the test statistic in the null distribution. Grey line highlights the value of 0.025 or 0.975. Difference lower than 0.025 represents the disparity of the former lower than the latter significantly, and the condition of the difference higher than 0.975 was inverse. Abbreviations: G, genus database; S, species database; pov, product of variance; sov, sum of variance; P1, Early Permian; P2, Middle Permian; P3, Late Permian; T1, Early Triassic; T2, Middle Triassic.

**Appendix figure 8.**
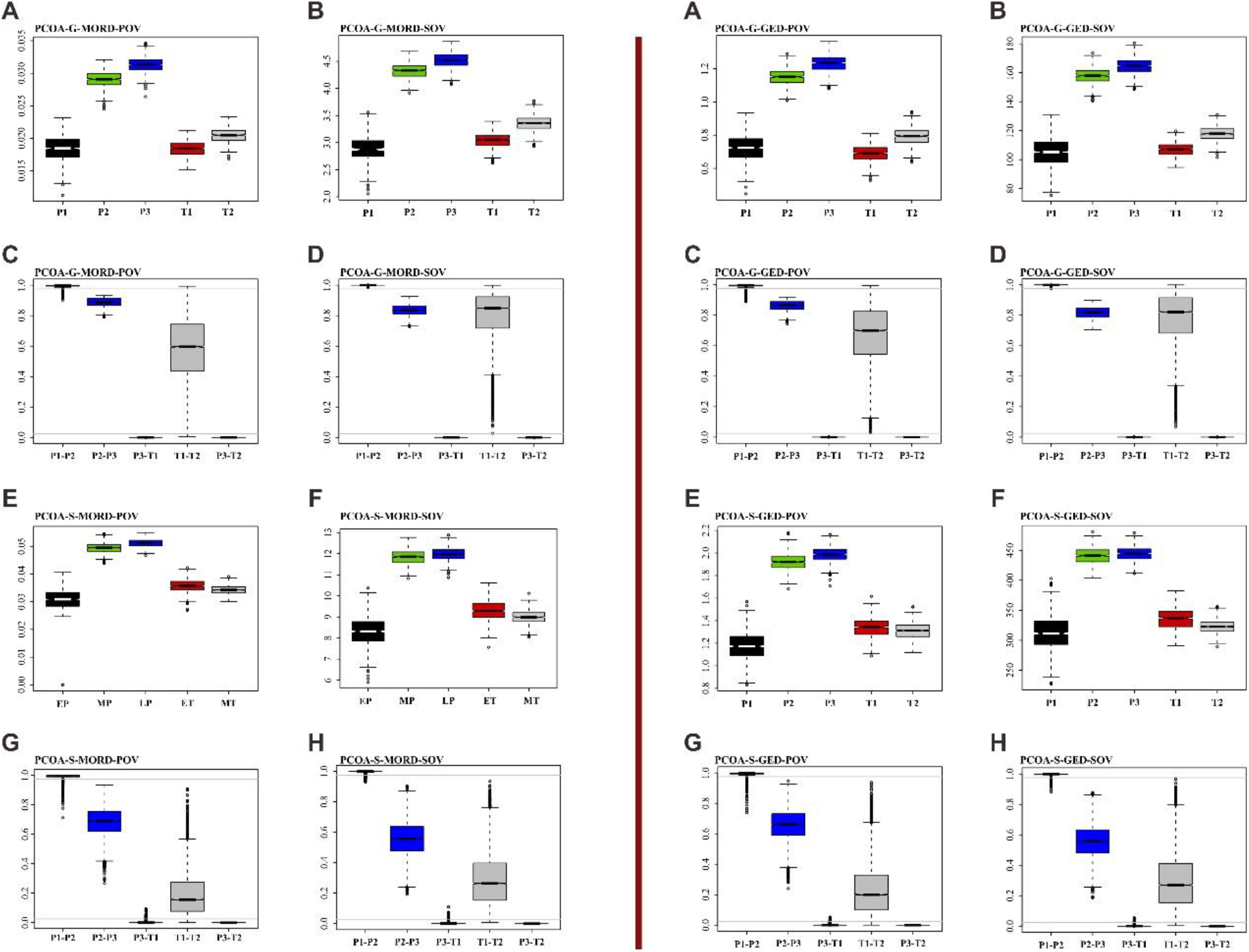
Disparity comparison and permutation tests with sample size corrected, under ordination method of PcoA, assuming that the age of the Grès à Voltzia specimens is Early Triassic. Left: results based on MORD matrix. Right: results based on GED matrix. **(A, B, E and F)** Disparity comparison. **(C, D, G and H)** Permutation tests with sample size orrected. Abbreviations: G, genus database; S, species database; pov, product of variance; sov, sum of variance; P1, Early Permian; P2, Middle Permian; P3, Late Permian; T1, Early Triassic; T2, Middle Triassic.

**Appendix figure 9.**
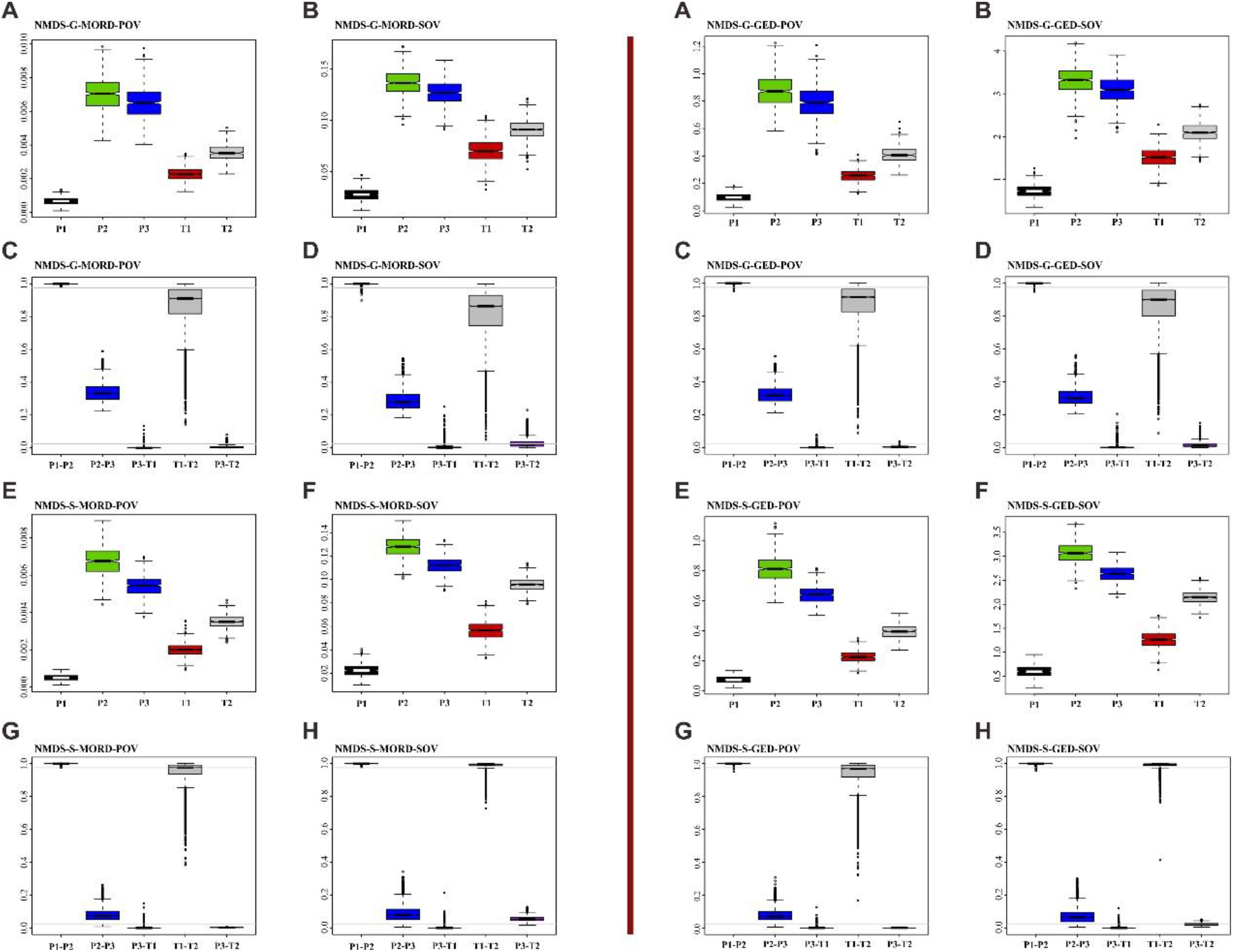
Disparity comparison and permutation tests with sample size corrected, under ordination method of NMDS, assuming that the age of the Grès à Voltzia specimens is Early Triassic. Left: results based on MORD matrix. Right: results based on GED matrix. **(A, B, E and F)** Disparity comparison. **(C, D, G and H)** Permutation tests with sample size corrected. Abbreviations: G, genus database; S, species database; pov, product of variance; sov, sum of variance; P1, Early Permian; P2, Middle Permian; P3, Late Permian; T1, Early Triassic; T2, Middle Triassic.

**Appendix figure 10.**
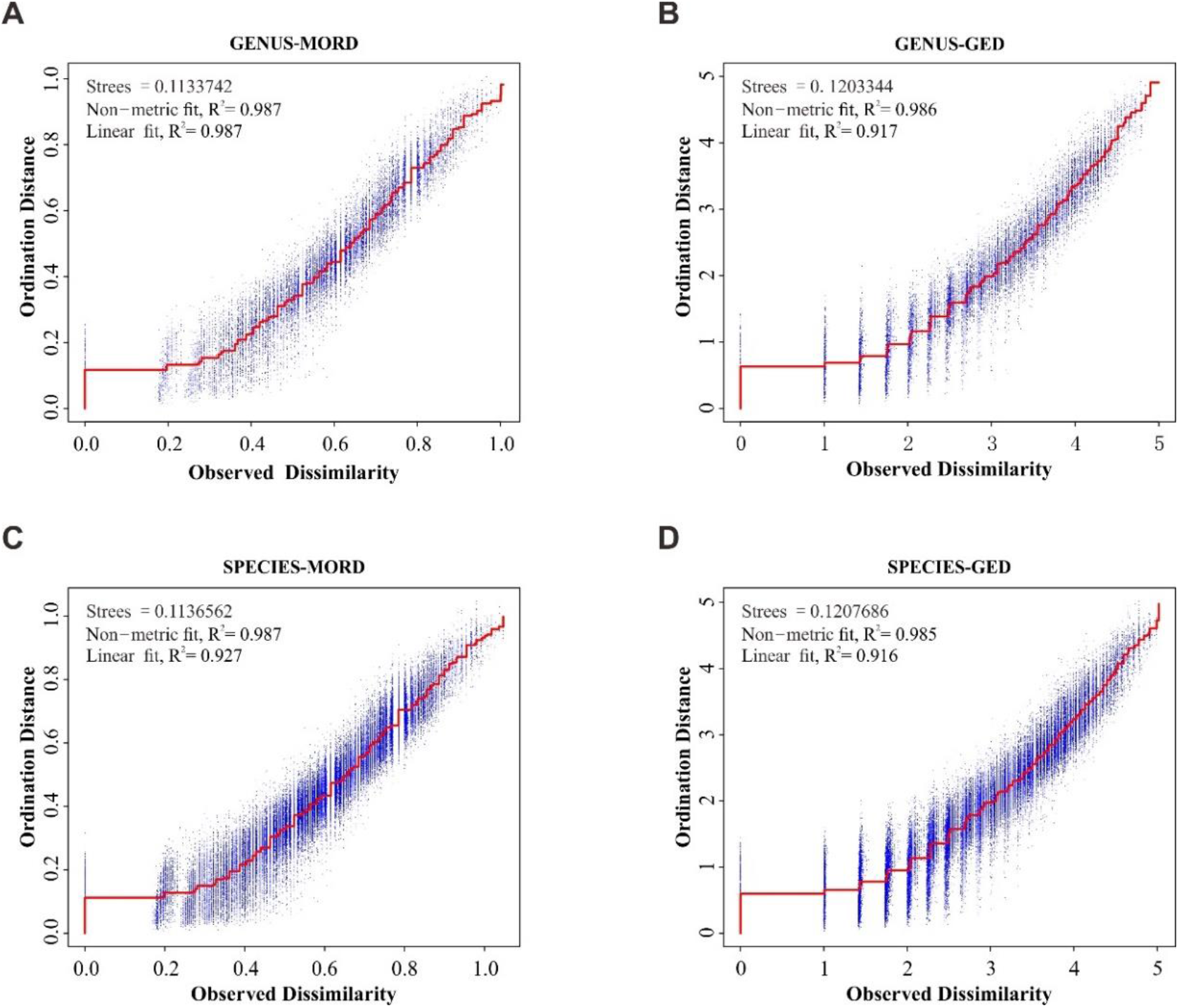
Shepard “goodness-of-fit” stress plot. (**A**) Result for MORD matrix of genus database. (**B**) Result for MORD matrix of species database. (**C**) Result for GED matrix of genus database. (**D**) Result for GED matrix of species database. Stress and R^2^ value indicate a good result of ordinations.

**Data S1.** (separate file) Fossil coleoptera database.

**Data S2.** (separate file) Character state matrix for the phylogenetic analysis.

**Data S3.** (separate file) Fossil character matrix for the morphospace analysis.

**Data S4.** (separate file) Result of permutation test for morphological disparity.

